# Biased enrichment of DNA uptake enhancing sequences in Pasteurellaceae and Neisseriaceae

**DOI:** 10.1101/2025.10.24.684340

**Authors:** Stian Aleksander Helsem, Kristian Alfsnes, Stephan A. Frye, Ole Herman Ambur, Alexander Hesselberg Løvestad

**Affiliations:** Department of Life Sciences and Health, Faculty of Health Sciences, OsloMet, Norway; Division for Infection Control, Norwegian Institute of Public Health, 0213 Oslo, Norway; Division of Laboratory Medicine, Dept. Microbiology, Oslo University Hospital, Norway

## Abstract

Some naturally transformable bacteria can selectively take up homologous DNA through short conserved motifs termed DNA Uptake Enhancing Sequences (DUES), comprising DNA Uptake Sequences (DUS) in *Neisseriaceae* and Uptake Signal Sequences (USS) in *Pasteurellaceae*. Using 177 complete genomes, this study provides the most extensive comparative analysis of DUES distribution, composition, and functional associations to date. Three novel DUS dialects were identified in *Neisseria animalis* and *Vitreoscilla* spp., extending the known diversity of the transformation system. Approximately half of all DUES and more than 90% of inverted repeat DUES occurred within predicted transcriptional terminators, and DUES inside coding-sequences were biased toward reading frames minimizing bioenergetic cost, indicating both structural and metabolic selection pressures. Gene Ontology and KEGG analyses revealed extensive but asymmetric functional enrichment: both families showed bias toward genome maintenance processes, yet *Neisseriaceae* displayed stronger enrichment for DNA repair, replication, and the UvrABC complex, whereas *Pasteurellaceae* were more associated with homologous recombination and the RecBCD complex. These patterns indicate that while DUES enrichment is broadly conserved, its functional integration diverges between families, reflecting distinct evolutionary adaptations that couple DNA uptake specificity to genome stability and cellular maintenance.

## Introduction

Bacterial transformation is the uptake and recombination of DNA from the environment, a process that promotes allelic reshuffling in ways reminiscent of meiosis in eukaryotes. Transformation is widespread in nature, and some of the enzymes involved are conserved across all domains of life. Bacteria have evolved diverse mechanisms for acquiring homologous DNA, but perhaps the clearest strategy for ensuring recombination within species boundaries is self-DNA recognition prior to uptake. In the two bacterial families Neisseriaceae and Pasteurellaceae, self-DNA recognition is mediated by short sequence motifs called DNA Uptake Sequences (DUSs) in the Neisseriaceae and Uptake Signal Sequences (USS) in the Pasteurellaceae (1–3). Here, we refer to these collectively as DNA Uptake Enhancing Sequences (DUES). DUESs are typically 9-10 nucleotides long, present in hundreds to thousands of copies in a genome, and appear completely unrelated between these two families (3). They may therefore represent two distinct but convergent adaptations to the same evolutionary need: securing homologous DNA for allelic replacement. However, recent findings by the authors have suggested that DUES may have evolved divergently (4). Within each family, several dialects of the uptake sequence have been described, with more closely related species often sharing a common sequence, while others exhibit similar but distinct variants (5,6). These dialects highlight the evolutionary flexibility of the system, maintaining the principle of self-recognition while restricting transformation primarily to within-species or within-genus boundaries. To date 8 dialects have been described in the Neisseriaceae and 2 in Pasteurellaceae. Recognition is mediated by specific proteins, ComP in Neisseria (7,8) and the recently predicted ComN in Pasteurellaceae (4), and has left a striking genomic imprint, with some species devoting up to ≈2% of their chromosome to DUES (6).

DUES can be located both within genes and in intergenic regions. Intergenic DUES often occur as inverted repeats that can form stem-loop structures functioning in transcriptional termination (5,9). Intragenic DUES, by contrast, may impose translational costs: they add amino acid constraints, and DUES in some reading frames encode stop codons, which must be avoided. As a result, the genomic distribution of DUES is shaped by negative positional constraints. Broadly, DUES tend to accumulate in permissive genomic regions, such as intergenic spaces or poorly conserved parts of genes, and they are biased toward certain functional categories of genes (10–12). Early genomic studies reported that DUES in both the Pasteurellaceae and Neisseriaceae families preferentially target genes involved in genome maintenance processes such as DNA repair, recombination, restriction-modification, and replication (10). These findings have been debated, with later studies questioning the strength and interpretation of such biases (11). A follow-up study using six neisserial genomes found that DUES were enriched in the core genome, depleted in regions under diversification, and absent from recently acquired or lost genes (12), suggesting a role for transformation in genome stability. However, these studies were based on limited genome datasets and broad functional categories, e.g. COG classifications, and many DUES-associated genes were annotated as hypothetical or of unknown function. As a result, the biological significance of DUES bias remains contested, with competing hypotheses on whether bacterial transformation primarily promotes adaptation, stability, or both (reviewed in (13)).

The rapid advancement of high-throughput sequencing technologies has led to an explosion of available bacterial genomes. This scale of data necessitates robust computational approaches for functional annotation, which in turn enable biological interpretation of genomic features such as uptake sequences. Functional annotation typically involves assigning Gene Ontology (GO) terms to genes, thereby translating raw sequence data into structured categories of function described in the GO knowledgebase (14). GO provides a hierarchical vocabulary covering three aspects of functional characteristics used to describe gene function: **Molecular Function (MF)** based on the activities performed by a gene product at the molecular level, **Biological Process (BP)** a “biological program” comprising molecular activities acting in concert to achieve a particular outcome; this program can be at the cellular level or at the organism level of multicellular organisms, and **Cellular Component (CC)** the locations, relative to cellular structures, where MFs are performed. GO terms are transferred to genes through orthology-based pipelines which are considered generally more accurate than similarity-only approaches such as BLAST or MMseqs2, since it minimizes incorrect functional transfer from paralogs (14). In parallel, Kyoto Encyclopedia of Genes and Genomes (KEGG) pathway annotation provides a complementary framework, mapping genes to metabolic and cellular pathways and thereby offering insight into how DUES may bias toward particular functional systems (15). Together, GO and KEGG resources enable systematic analyses of functional biases across large genomic datasets.

The objective of this study is therefore to perform a comprehensive comparative analysis of DUES across all currently available 177 high-quality genomes from the Neisseriaceae and Pasteurellaceae, including several recently discovered species with uncharacterized DUES profiles and dialects.

## Materials and methods

### Genome collection and DUES scanning

All Neisseriales and Pasteurellales genomes available at NCBI as of 24th of January 2025 were downloaded in nucleotide/protein FASTA and GFF3 format. The frequency of all 9- and 10-mers across all genomes were mapped and overrepresentation statistics were calculated using the approach described in (4), with a particular emphasis on DUES previously identified (6). To identify potentially novel DUES, known DUES were removed and candidate DUES trimmed 1-2 nt at the 5’- and 3’-end in order to avoid sequences overlapping with known DUES. Sequence logos of potential DUES dialect conservation were generated for their respective genomes, allowing for one mutation in the DUES sequence. We generated an alignment of the known and potentially new DUS dialects using MAFFT v7.505 (16). and visualized the alignment in MView v. 1.68 (17).

### Gene annotations

eggNOG-mapper v. 2.1.12 (18,19) were used to annotate CDS with KEGG and GO terms using the program parameters -m diamond, --itype proteins, --tax_scope none, --tax_scope_mode broadest, --sensmode ultra-sensitive. --decorate_gff and --go_evidence all.

### DUES reading frame usage

The reading frame bias of DUES residing in CDS were investigated to test the following hypothesis: (1) on average, energetically cheap reading frames are used more often than expensive ones in terms of amino acid biosynthetic ATP usage, and (2) that the six reading frames of DUES are on average cheaper than those of other 9-mers or 10-mers. To test this, the usage of the 20 different amino acids with regard to the amount of ATPs consumed in their biosynthesis was analyzed following the calculations by (20). We then (i) mapped the peptides resulting from all DUES occurrences across genomes, (ii) stored the associated ATP-cost and (iii) generated 1000 permutations from these data points, each time randomly drawing one of the six possible reading frames. The observed mean to the permuted mean was then compared using both a lower-tail (H_a_: observed mean is lower than permuted means; null hypothesis H_0_: observed mean is not different from permuted means) and two-sided (alternative hypothesis H_a_: observed mean is higher or lower than permuted means) empirical p-value.

Additionally, to test whether the peptides DUES contribute to on average are cheaper than those contributed by non-DUES of similar length (H_0_: No difference in average reading frame costs of DUES and permutations), we permuted the DUES genomic coordinates (100 permutations per species) using different 9- and 10-mer motifs for each original DUES occurrence. The ATP-cost of both original DUES contributions and contributions by permuted motifs were then calculated and the means compared using Welch’s t-test.

### DUES transcriptional terminators

Transcriptional terminators were predicted using BacTermFinder v. 1.0 (21), using default parameters. Overlapping coordinates of predicted terminators and DUES coordinates were found using BEDTools v2.31.1 (22). Additionally, we annotated the DUES that qualified as inverted repeat DUES (irDUES), following the definition in Spencer-Smith et al. 2016 (9).

### GO Enrichment and KEGG pathway Analysis

Genes containing DUES motifs were identified, and their collected associated GO terms were tested for over-representation in 107 species. Of these, 49 species were from the Pasteurellaceae genera *Actinobacillus* (N=15), *Aggregatibacter* (N=4), *Haemophilus* (N=11), *Mannheimia* (N=10), *Pasteurella* (N=9) and 58 species from the *Neisseriaceae* genera *Alysiella* (N=2), *Eikenella* (N=5), *Kingella* (N=7), *Vitreoscilla* (N=3) and *Neisseria* (N=41). The enrichment analysis was performed separately for the BP, MF, and CC ontologies using the R package topGO (23). The `weight01` algorithm, which incorporates GO hierarchy information, was used with Fisher’s Exact Test to calculate significance against a background of all annotated genes in each genome. GO terms with a p-value < 0.05 in the weight01-analysis were considered significant. Following the topGO authors’ recommendation from the manual, a multiple testing correction was not applied, as the chosen algorithm is designed to address the issue of GO term dependency and reported p-values should be considered already corrected.

KEGG pathway-enrichment analysis was performed on KEGG Orthology (KO) term assignments from EggNOG-mapper. For each species, all genes overlapping a US motif formed the test set and all KO-annotated genes in that genome formed the background. KO→pathway mappings were retrieved via KEGGREST (24) and tested with clusterProfiler’s enricher (25). Significance was defined as Benjamini-Hochberg-adjusted p < 0.10.

## Results

### DUES scanning reveals three novel DUES dialects in Neisseriaceae

The DUES scanning revealed three potentially novel DUS variants. In *Neisseria animalis* strain ATCC 49930, the 10-mers **GCCGTCTGCA** (2,296 copies; observed/expected ratio = 15.7; adjusted *p* = 0) and the overlapping **CGTCTGCAAA** (1,813 copies; ratio = 27.3; adjusted *p* = 0) were highly overrepresented. Together, these sequences define the 12-mer DUS dialect **GCCGTCTGCAAA**, which appears strongly conserved in this strain (Fig. 1 A). Relative to the canonical DUS **GCCGTCTGAA** documented in many *Neisseria* species including *N. gonorrhoeae* and *N. meningitidis* (6) the dialect found in *N. animalis* differs only by one A-C transversion variation. As shown in Fig 1 A, this transversion variation is also the least conserved relative to all other positions of the DUES.

**Figure 1.**
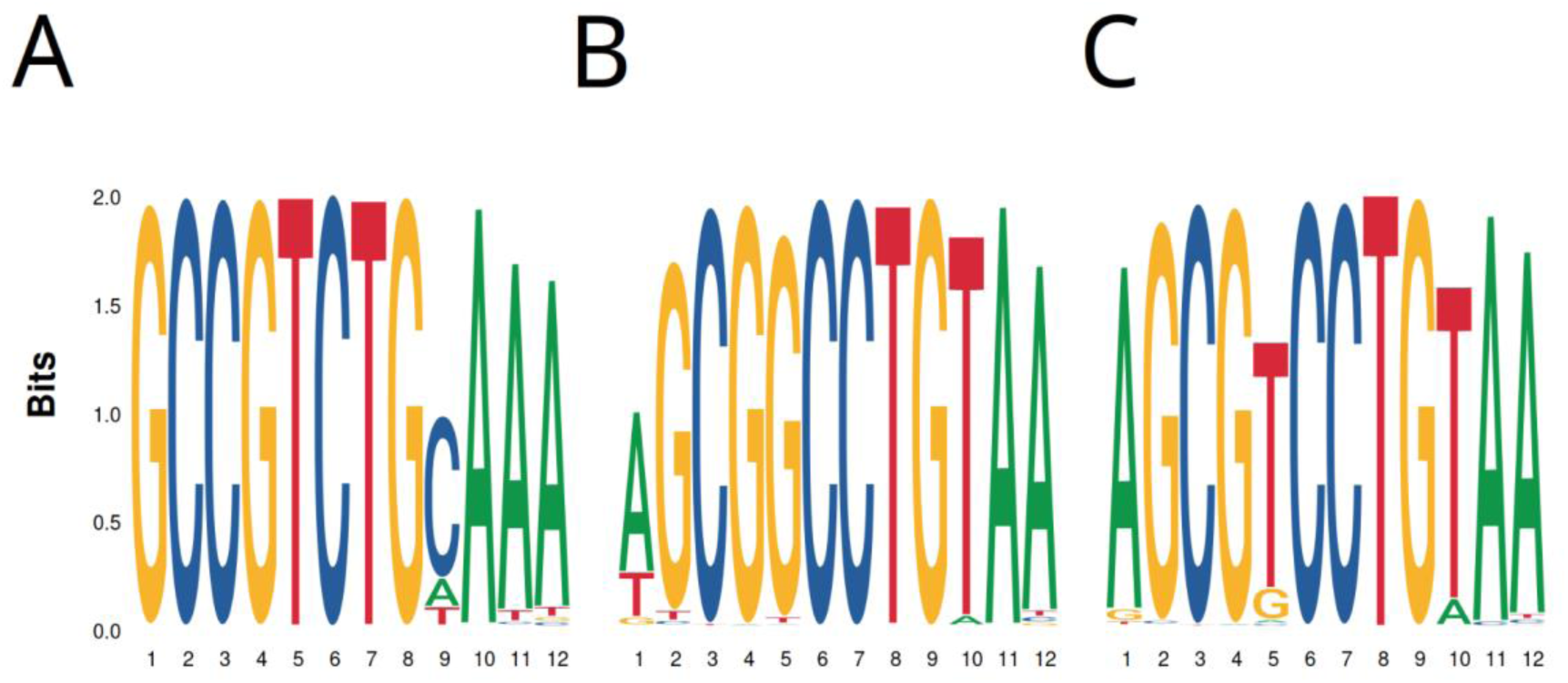
Genomic DUS conservation. Genomic sequence logos of the novel DUS dialects identified in (A) *Neisseria animalis* strain ATCC 49930, (B) *Vitreoscilla massiliensis* strain SN6, shown with one mismatch allowed and (C) *Vitreoscilla stercoraria* strain SAG 1488-6

In *Vitreoscilla stercoraria* strain SAG 1488-6, the 10-mers **CGTCCTGTAA** (1,291 copies; ratio = 49.3; adjusted *p* = 0) and the overlapping **AGCGTCCTGT** (1,234 copies; ratio = 45.8; adjusted *p* = 0) specify the proposed 12-mer dialect **AGCGTCCTGTAA** (Fig. 1 B). A variant of this motif (**AGCGGCCTGTAA**) was detected in *Vitreoscilla* massiliensis strain SN6, differing by a T→G transversion variation relative to the *V. stercoraria* sequence. The overlapping 10-mers **CGGCCTGTAA** (3,105 copies; ratio = 65.0) and **AGCGGCCTGT** (2,129 copies; ratio = 47.9) support this interpretation (Fig. 1 C). Relative to the canonical *Neisseria* DUS **GCCGTCTGAA,** the novel dialects identified in the Vitreoscillae, which both contain the functionally relevant CTG core, are characterized by two transversion variation C-G and A-T, one transition variation T-C; and the *V. massiliensis* dialect an additional G-T transversion variation relative to both the canonical and *V. stercoraria* dialects (Fig. 2).

**Figure 2.**
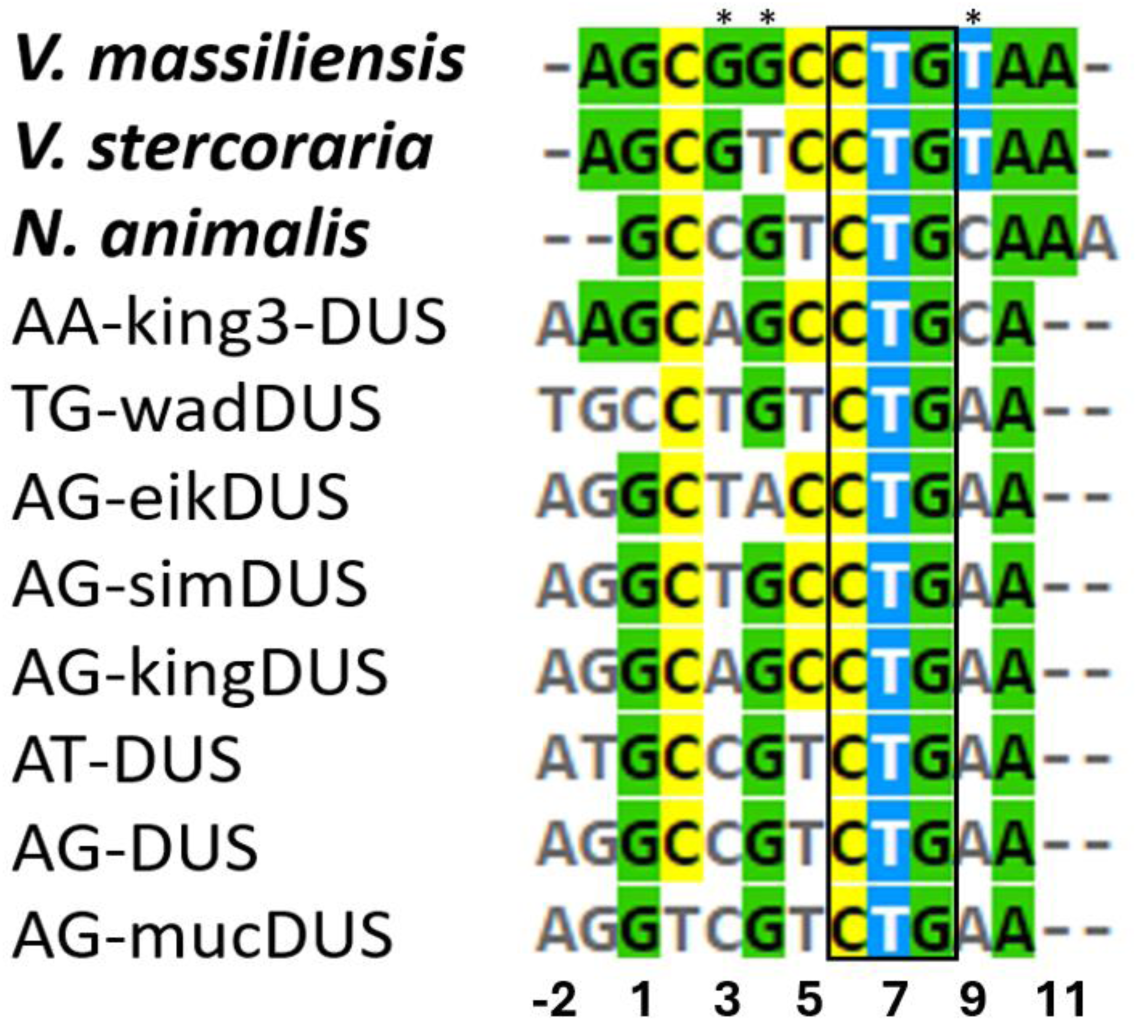
Alignment of different DUS dialects. The three new identified DUS dialects are labelled by species name in bold above previously assigned dialects. The dialect unique nucleotides observed in the two *Vitreoscilla* species are labelled with an asterix. The CTG core, previously shown essential for transformation in *Neisseria* (6), is boxed. The coloring follows the consensus for the particular position and numbering to previously established convention with CTG core in positions 6-8 (6).

### US reading frames are biased towards lower biosynthetic energy cost

Analysis of ATP-costs showed that DUES are preferentially positioned in coding sequences to reduce energetic burden (Fig. 3). Median and mean frame costs were lower for observed DUES compared to permuted controls (−1 vs. 0; −0.6 vs. 2.14), and the difference was highly significant (Welch’s *t*-test, *p* < 2.2 × 10⁻¹⁶). Similarly, permutation analysis of genomic coordinates revealed that the average cost of DUES was significantly lower than that of general 9- and 10-mers (∼0.39 vs. 1.15; *p* = 2.8 × 10⁻⁶).

**Figure 3.**
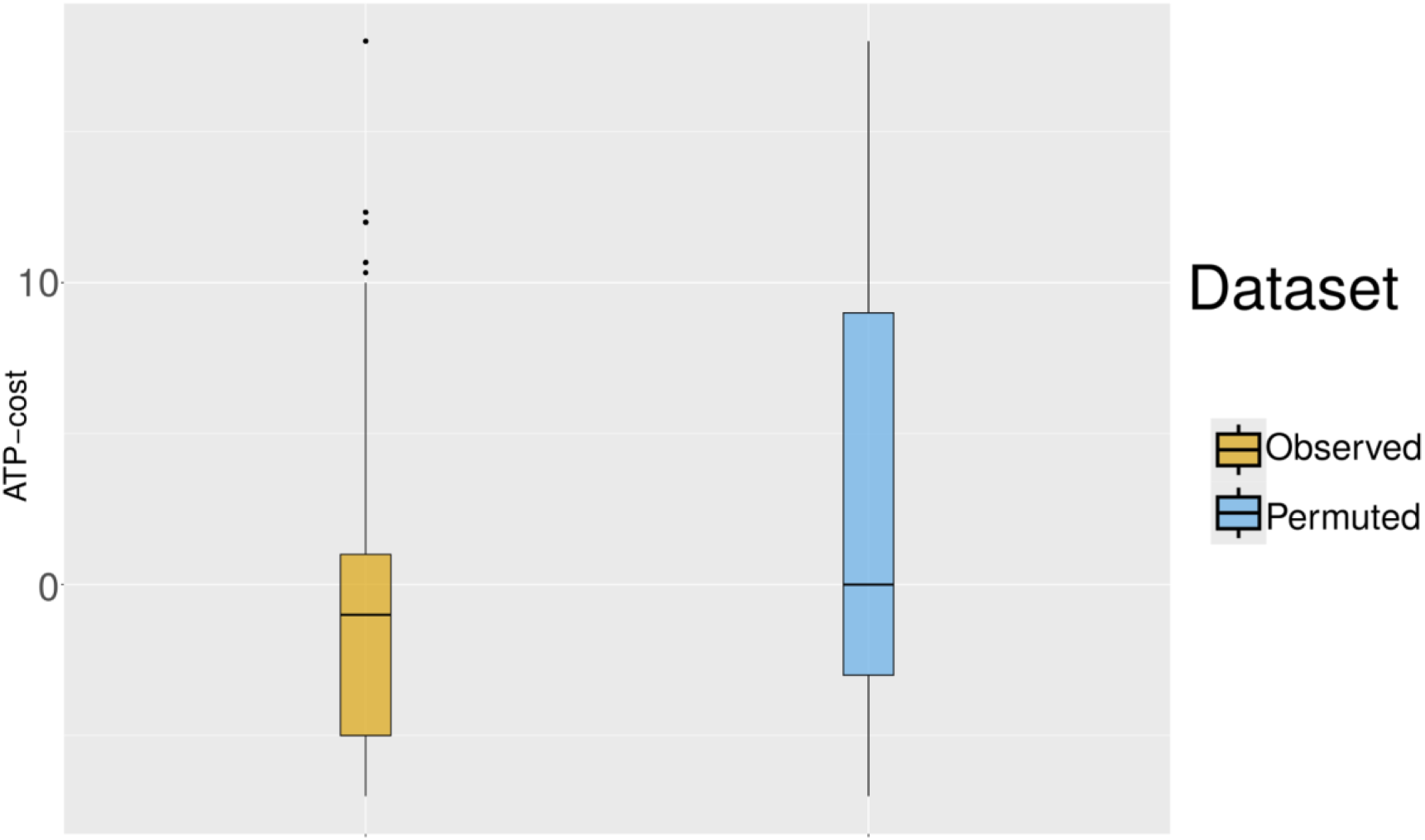
Box plots showing the distribution of ATP-costs for observed reading frames and 1000 permuted sets across all species and DUES, showing a clear skew toward cheaper reading frames among actual DUES relative to permuted ones. The median is shown as horizontal bars inside boxes, which depict interquartile ranges (IQR), and the whiskers extend 1.5 the IQR.

Frame usage analysis across species revealed distinct biases for each dialect (Fig. 5). In Pasteurellaceae, **Hin-USS**, TAL and SAV were the most frequent frames, while KCG was least common. In Pasteurellaceae, **Apl-USS** was dominated by QAV (∼50% of cases), with KRX least frequent. The Neisserial **canonical DUS** motifs showed a fairly even distribution across four frames, with SDG slightly enriched, while AV*X (* = stop codon) and FRRX (X = any codon) were rarely seen. These findings are in line with previous research (11). In **wadDUS**, LSX and ACLX predominated, and PV* was least common. **eikDUS** exhibited roughly equal distribution among four frames, although FR*X was least frequent. In **mucDUS**, RLX and SDD occurred at similar frequencies, while FRRX and VV*X were infrequent. **kingDUS** was dominated by SLX and QPE, with SGC the least used. **king3DUS** showed enrichment of SLX and QAA, with CRLX rare. **simDUS** displayed comparable frequencies of LPE and SGS, with FRQX least common. Newly described dialects also exhibited strong frame biases: **vitDUS** was dominated by QDA (>50%), **aniDUS** by ADG (>40%), and **vit3DUS** by QAA (>50%).

Together, these patterns demonstrate that DUES across both already established and novel DUS dialects are not randomly distributed among frames but consistently favor a subset of energetically less costly codon contexts, reinforcing selective pressures on their genomic positioning.

### Frequent localization of DUES within predicted terminators

The fraction of uptake signal sequences located within predicted transcriptional terminators was similar between the two families. On average, about half of all DUES in Neisseriaceae and Pasteurellaceae were positioned inside BacTermFinder-predicted terminators (Fig. 5). This proportion is notably higher than reported in earlier studies based on smaller datasets, which estimated that only 16-29% (9,103) or 13-21% (9,103) of predicted terminators contained DUES.

Inverted-repeat DUES showed an even stronger tendency to occur within terminators, particularly in Pasteurellaceae, contrasting with earlier findings based on GeSTer (9,104,105) and TransTermHP (103) predictions (Figs. 4.3.4-4.3.5). The mean and median fractions of irDUES within terminators in Pasteurellaceae were ∼0.88 and ∼0.91, respectively, compared to ∼0.70 and ∼0.74 in Neisseriaceae.

**Figure 4.**
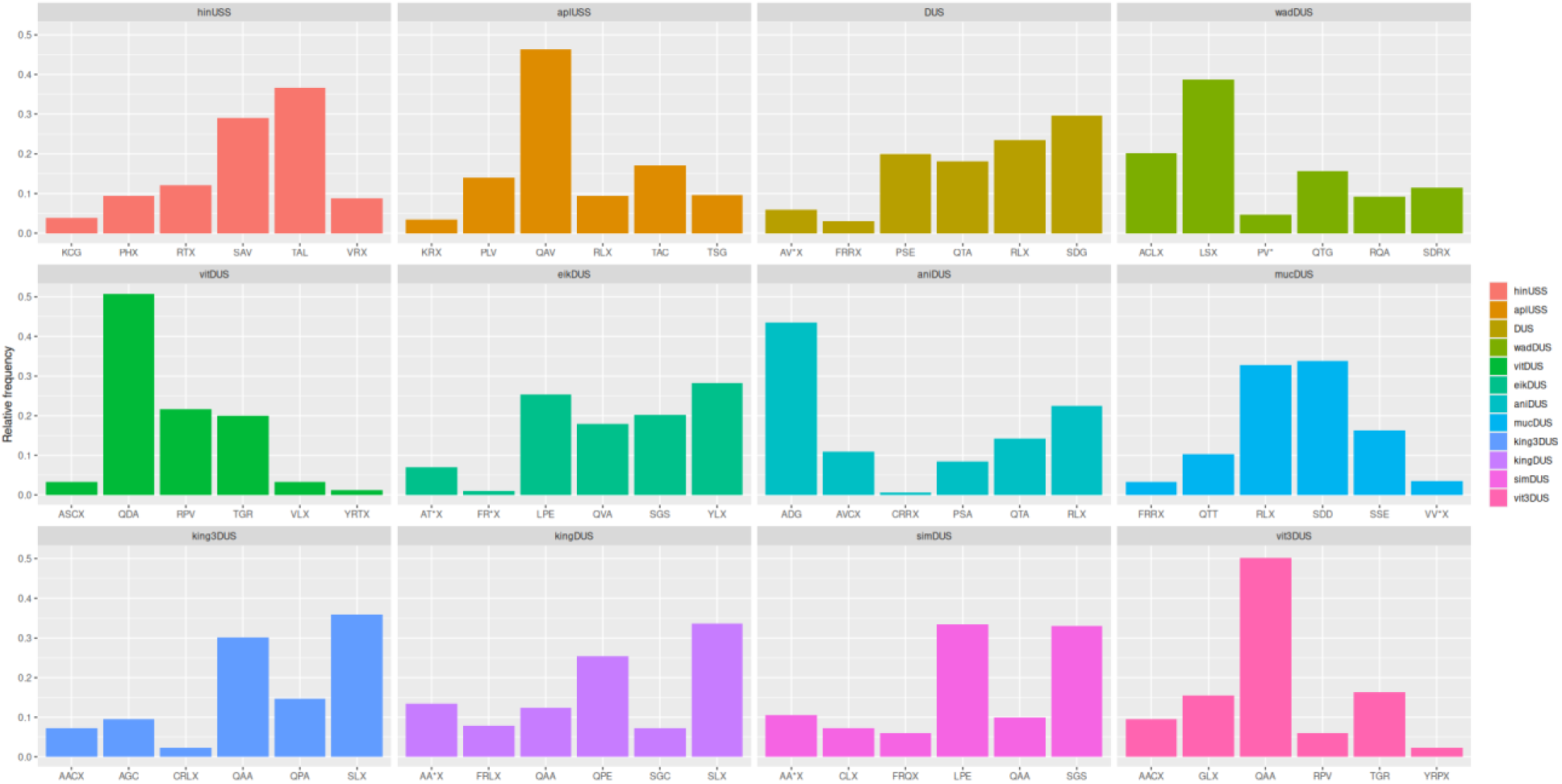
Observed relative frequencies of the possible reading frames per DUES. * = stop codon. X = Ambiguous amino acid.

Family-specific trends were also observed when considering species with higher motif counts. In Neisseriaceae species with ≥1000 total DUES, the median and mean fractions of the dominant dialect located within terminators were 1.0 and 0.91, respectively. In Pasteurellaceae species with ≥400 total DUES, the corresponding values were 0.96 and 0.94. These distributions closely mirrored those obtained for the overall fractions of DUES in terminators, as well as their dominant dialect equivalents (Fig. 5). Comprehensive statistics for all terminator prediction analyses are provided in Supplementary Tables 1 and 2.

**Figure 5.**
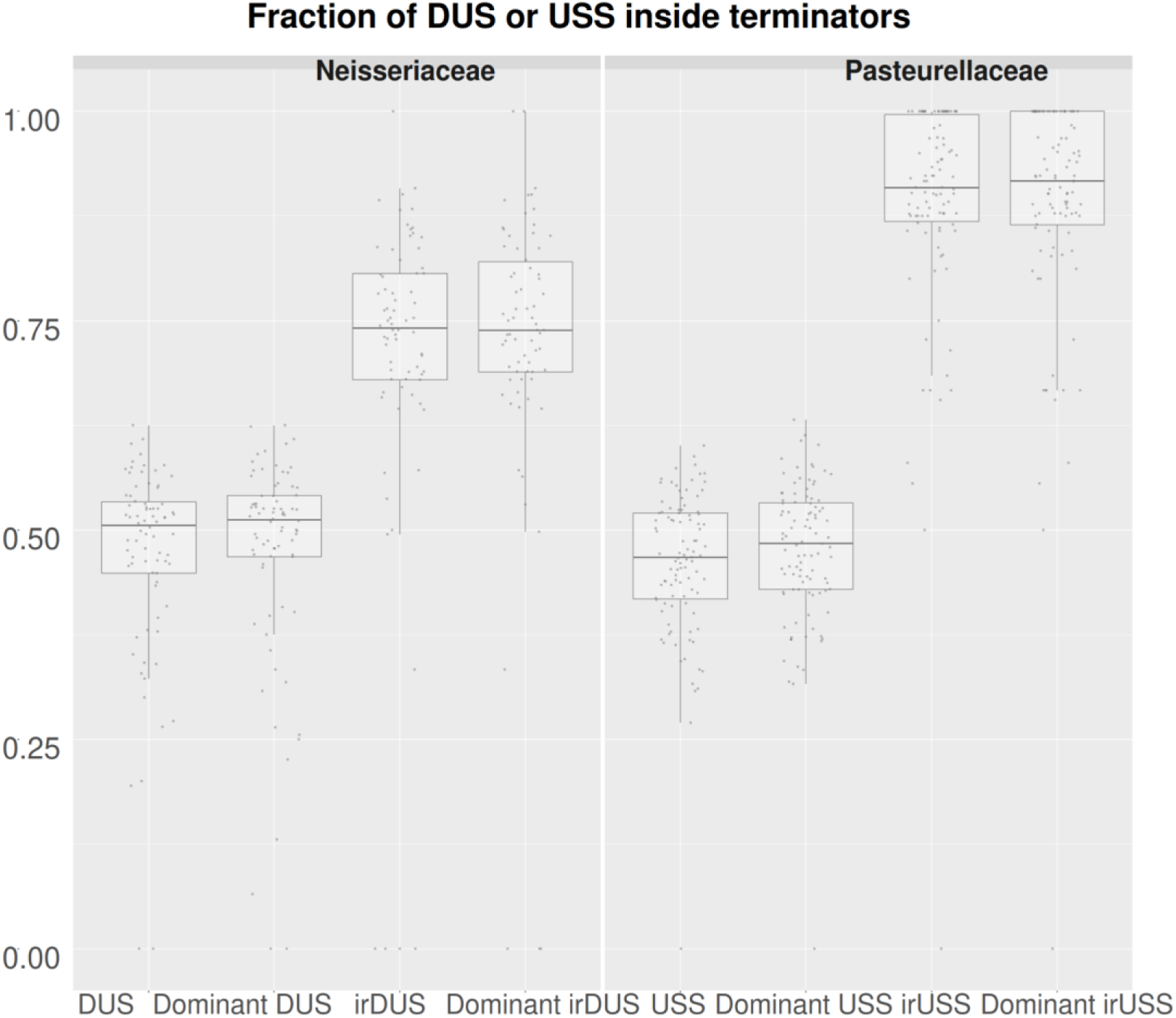
Box plots showing the distribution of DUS_term_/DUS_total_ and USS_term_/USS_total_ across all species.

### Overview of detected and enriched GO terms in Pasteurellaceae and Neisseriaceae

Across Pasteurellaceae and Neisseriaceae, 3646, 2317 and 355unique GO terms (i.e. GO terms found multiple times within each family are only reported once) were identified from the BP, MF and CC ontologies, respectively (Fig. 6). Of these GO terms, 67.7% BP GO terms, 71.2% MF GO terms and 70.1% CC GO terms were found in both families. 2750 BP GO terms, 1913 MF GO terms and 298 CC GO terms were assigned to genes containing at least one DUES motif, and 66.3%, 65.6% and 62.4% of these were shared across the two families, respectively. 266 BP GO terms, 230 MF GO terms and 52 CC GO terms were enriched in Pasteurellaceae and/or Neisseriaceae, with 24.4%, 21.7%, and 34.6% of them being enriched in both families, respectively. In total, 132 GO terms were enriched in both Neisseriaceae and Pasteurellaceae (65 BP, 50 MF and 18 CC).

**Figure 6.**
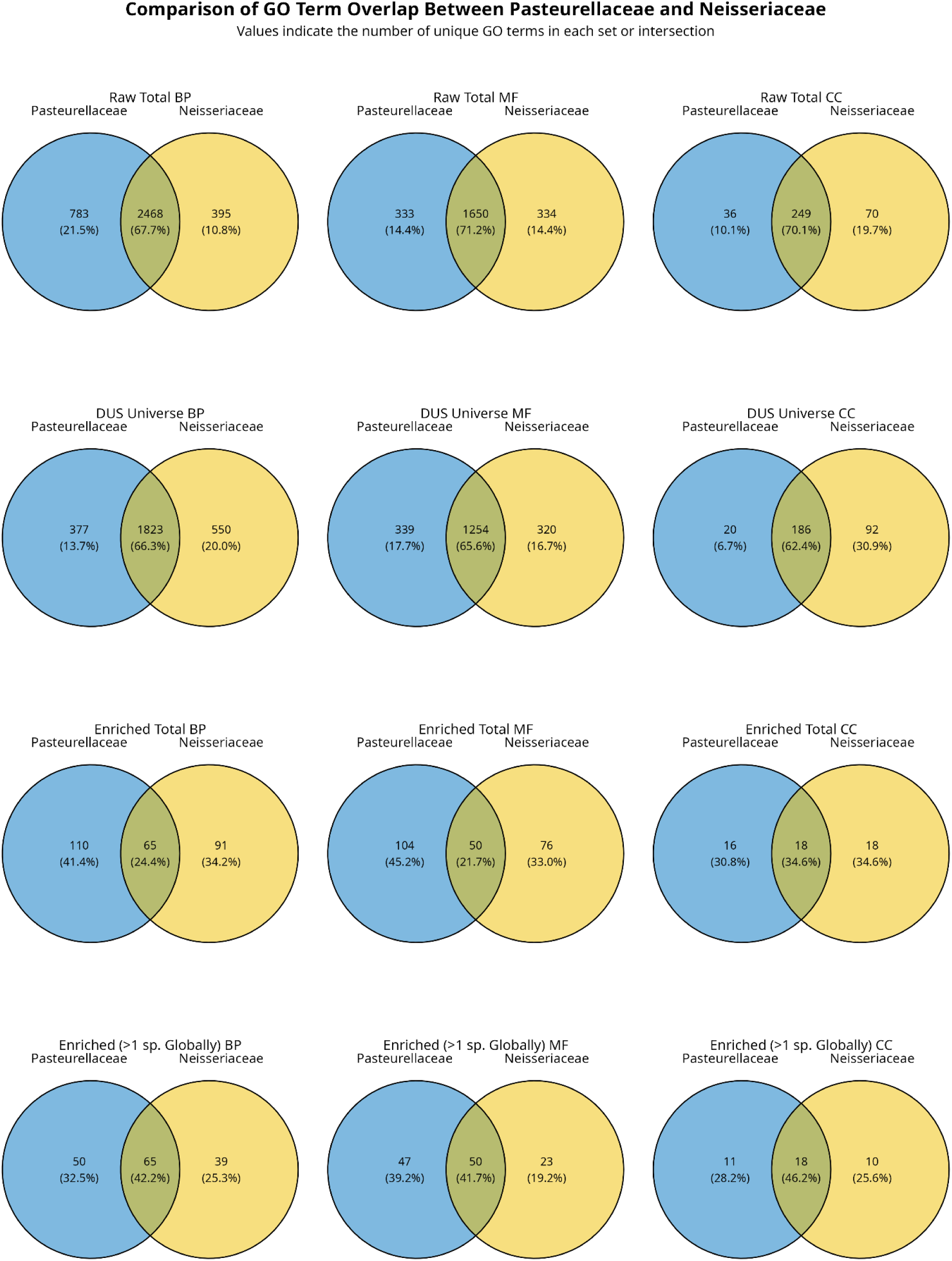
Venn diagrams showing Gene Ontology (GO) term overlap between Pasteurellaceae (blue, left) and Neisseriaceae (yellow, right) (circles not in scale). Each panel indicates the number of unique GO terms specific to each family or shared between them. The first row shows all GO terms identified across species within each family. The second row shows GO terms associated with genes containing DUES. The third row shows enriched GO terms within each family, while the fourth row shows enriched GO terms found in more than one species.

Many enriched terms were restricted to a single species, ranging from 31.3% to 69.8% across all ontologies. The CC ontology showed the lowest proportion of species-unique terms (31.3% in Pasteurellaceae, 44.5% in Neisseriaceae), whereas BP and MF ontologies had much higher proportions of species-unique terms (BP: 54.5% in Pasteurellaceae, 57.2% in Neisseriaceae; MF: 54.9% in Pasteurellaceae, 69.8% in Neisseriaceae).

### BP GO terms

The BP GO term “DNA geometric change” was enriched in more than 40% of the species in both Neisseriaceae (82.8%) and Pasteurellaceae (53.1%), while for the other shared terms Neisseriaceae showed higher enrichment than Pasteurellaceae (≥15%) for a broad set of functions (Fig. 7). These included “carboxylic acid metabolic process” (94.8% vs. 6.1%), “cell division” (34.5% vs. 2.0%), “cell wall macromolecule metabolism” (46.6% vs. 6.1%), “cell wall organization” (39.7% vs. 4.1%), “DNA biosynthetic process” (67.2% vs. 36.7%), “DNA replication” (50.0% vs. 12.2%), “DNA-templated DNA replication” (31.0% vs. 10.2%), “macromolecule biosynthetic process” (44.8% vs. 2.0%), “mismatch repair” (32.8% vs. 2.0%), “nucleotide-excision repair” (39.7% vs. 16.3%), “peptidoglycan biosynthesis” (48.3% vs. 14.3%), “regulation of cell shape” (46.6% vs. 6.1%), “response to radiation” (31.0% vs. 10.2%), “RNA modification” (77.6% vs. 4.1%), and “rRNA base methylation” (39.7% vs. 16.3%).

**Figure 7.**
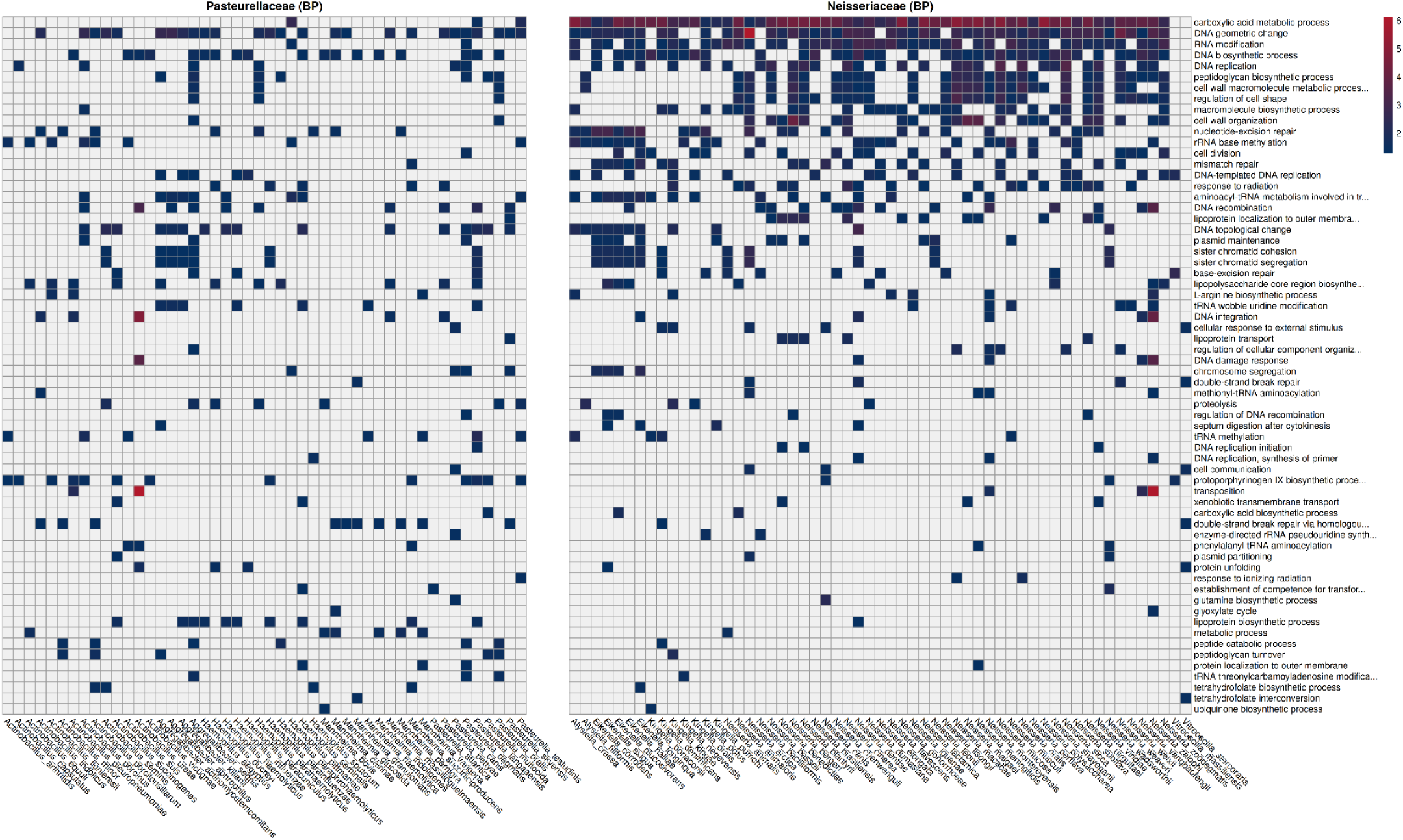
Enriched BP GO terms shared between Pasteurellaceae and Neisseriaceae. GO-terms are sorted by proportion of enriched species in the different families, with the most enriched GO term at the top. Color scale denotes -log10 adjusted p-values.

Pasteurellaceae displayed fewer strongly biased terms, with only “double-strand break repair via homologous recombination” (20.4% vs. 3.4%), “lipoprotein biosynthetic process” (20.4% vs. 1.7%), and “protoporphyrinogen IX biosynthesis” (26.5% vs. 5.2%) having ≥15% higher enrichment than in Neisseriaceae.

Additionally, some enriched BP GO terms were unique to each family. In Neisseriaceae, 31% of species were enriched for “DNA repair” and 14% for “SOS response,” whereas in Pasteurellaceae 31% of species were enriched for “tRNA processing” (Supplementary Table X). Together, these patterns indicate that while both families share a common set of BP terms, Neisseriaceae species show systematically higher enrichment across processes related to genome maintenance, DNA repair, and basic cellular functions, whereas Pasteurellaceae displayed narrower but distinct enrichments in homologous recombination (also a DNA repair mechanism) and metabolic specializations.

### MF GO terms

Several MF GO terms were enriched in more than 40% of the species in both Neisseriaceae and Pasteurellaceae. These included “ATP binding” (98.3% in Neisseriaceae vs. 98.0% in Pasteurellaceae), “ATP hydrolysis activity” (94.8% vs. 87.8%), “DNA binding” (77.6% vs. 46.9%), “DNA-directed DNA polymerase activity” (72.4% vs. 53.1%), “DNA helicase activity” (70.7% vs. 46.9%), and “exodeoxyribonuclease V activity” (44.8% vs. 46.9%) (Fig. 8).

**Figure 8.**
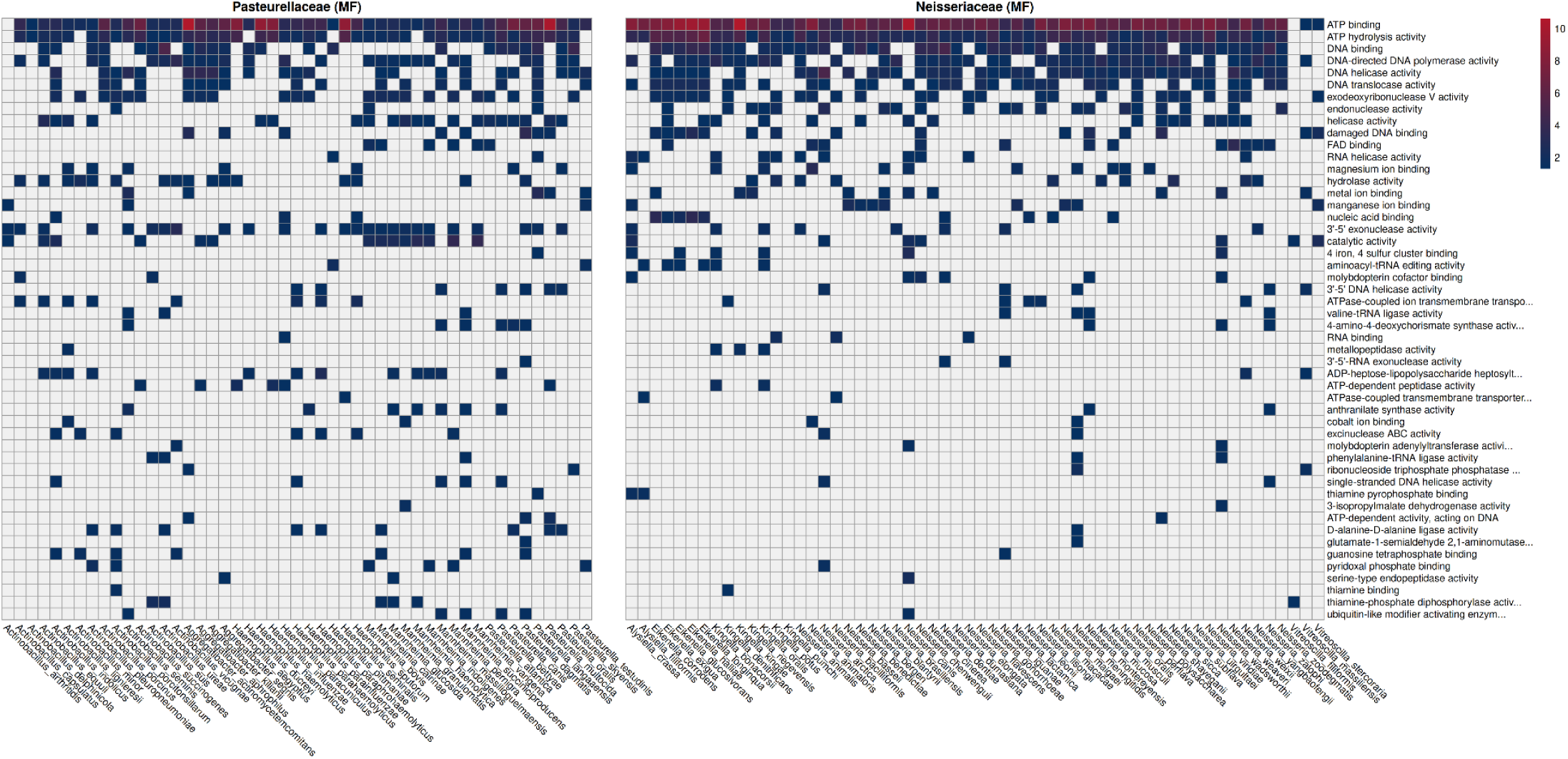
Enriched MF GO terms shared between Pasteurellaceae and Neisseriaceae. GO-terms are sorted by proportion of enriched species in the different families, with the most enriched GO term at the top. Color scale denotes -log10 adjusted p-values.

Neisseriaceae showed stronger enrichment than Pasteurellaceae (≥15% difference) for several DNA-associated functions. These included the aforementioned “DNA binding”, “DNA-directed DNA polymerase activity” and “DNA helicase activity” in addition to “DNA translocase activity” (58.6% vs. 34.7%), and “endonuclease activity” (36.2% vs. 6.1%).

Pasteurellaceae exhibited stronger enrichment for a smaller set of molecular functions. These included “hydrolase activity” (32.7% vs. 17.2%), “3′-5′ exonuclease activity” (51.0% vs. 13.8%), “catalytic activity” (28.6% vs. 12.1%), “ADP-heptose-lipopolysaccharide heptosyltransferase activity” (24.5% vs. 3.4%), and “D-alanine-D-alanine ligase activity” (18.4% vs. 1.7%). Additionally, Pasteurellaceae species were uniquely enriched for the terms “exoribonuclease II activity” (29%) and “DNA topoisomerase type I (single strand cut, ATP-independent) activity” (27%).

Overall, both families shared a strong enrichment in core enzymatic, ATP and DNA-associated terms, but Neisseriaceae displayed a more pronounced bias toward DNA-associated terms, while Pasteurellaceae exhibited stronger enrichment in metabolic and catalytic processes related to cell wall and lipopolysaccharide biosynthesis.

### CC GO terms

No CC GO terms were enriched in more than 40% of the species in both Neisseriaceae and Pasteurellaceae. Two terms showed stronger signals of enrichment in Neisseriaceae species (≥15% difference). These included “DNA topoisomerase type II (double strand cut, ATP-hydrolyzing) complex” (17.3% in Neisseriaceae vs. 2.1% in Pasteurellaceae) and “excinuclease repair complex” (73.1% vs. 8.3%). Neisseriaceae species were also solely enriched in the GO terms “DNA repair complex” (30.7%) and “mismatch repair complex” (26.9%).

Pasteurellaceae showed stronger enrichment in four CC GO terms. These included “DNA polymerase III core complex” (37.5% in Pasteurellaceae vs. 9.6% in Neisseriaceae), “cytoplasm” (45.8% vs. 15.4%), “exodeoxyribonuclease V complex” (58.3% vs. 17.3%), and “plasma membrane-derived chromatophore membrane” (25.0% vs. 1.9%).

Overall, there were little overlap between the families, but they both show a biased enrichment for separate DNA repair complexes, where Neisseriaceae had a high enrichment for excinuclease repair complex (UvrABC) and Pasteurellaceae was more enriched for exodeoxyribonuclease V complex (RecBCD).

**Figure 9.**
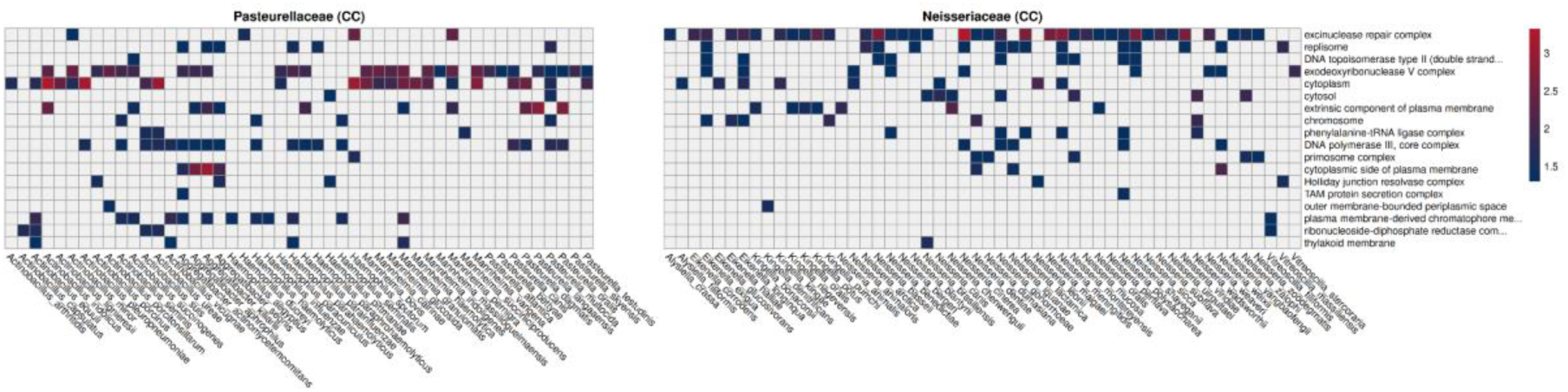
Enriched CC GO terms shared between Pasteurellaceae and Neisseriaceae. GO-terms are sorted by proportion of enriched species in the different families, with the most enriched GO term at the top. Color scale denotes -log10 adjusted p-values.

### KEGG pathway analysis

The KEGG pathway revealed a high enrichment for “Aminoacyl-tRNA biosynthesis pathways” as well as several species enriched for the genome maintenance-related pathways “Nucleotide excision repair”, “Mismatch repair” and “Homologous recombination” in Neisseriaceae (Fig. 10). Shared enrichment was not found in any particular pathway in the investigated Pasteurellaceae species (Fig. 11).

**Figure 10.**
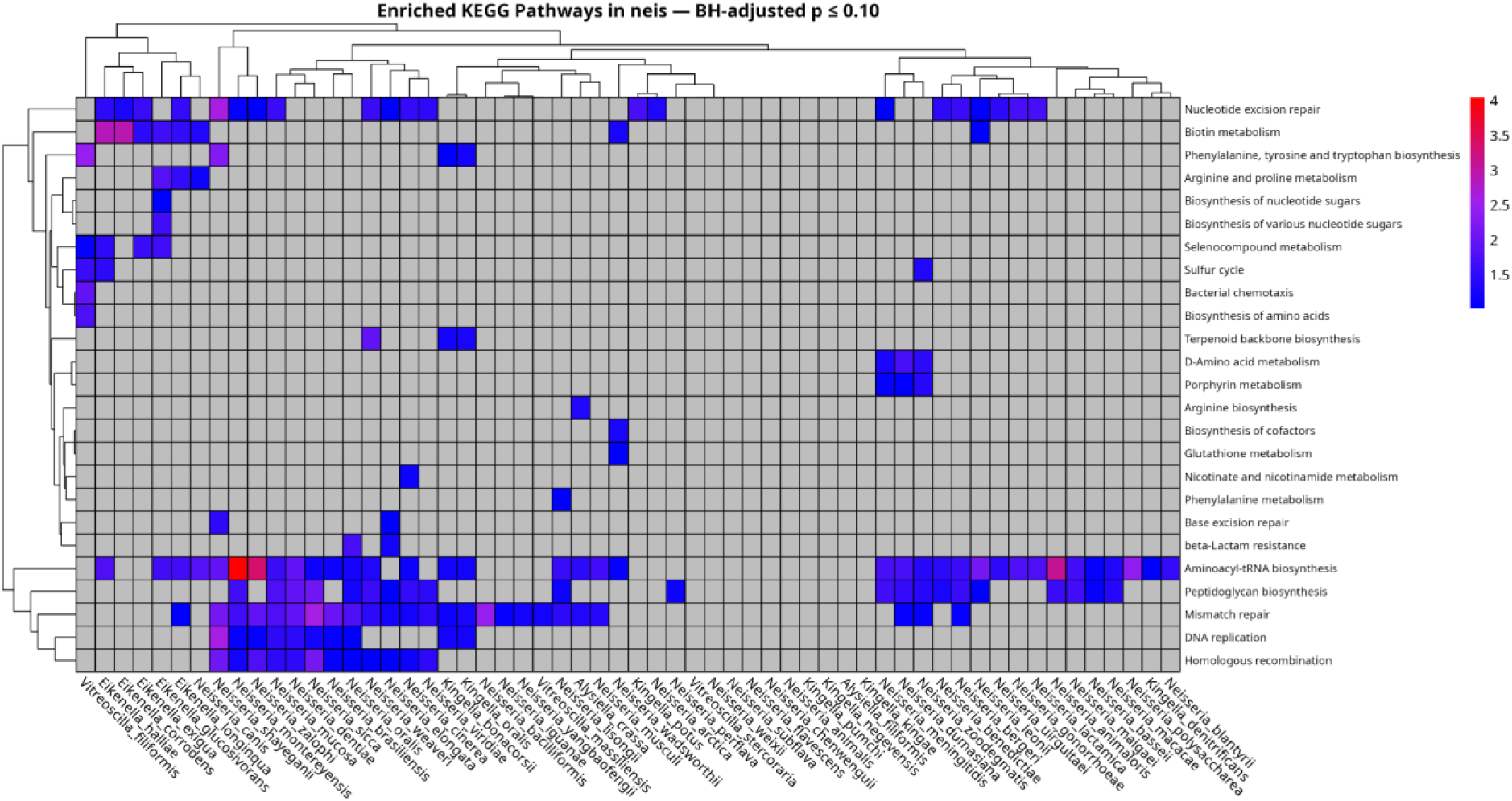
Enriched KEGG pathways in Neisseriaceae. Color scale denotes -log10 adjusted p-values.

**Figure 11.**
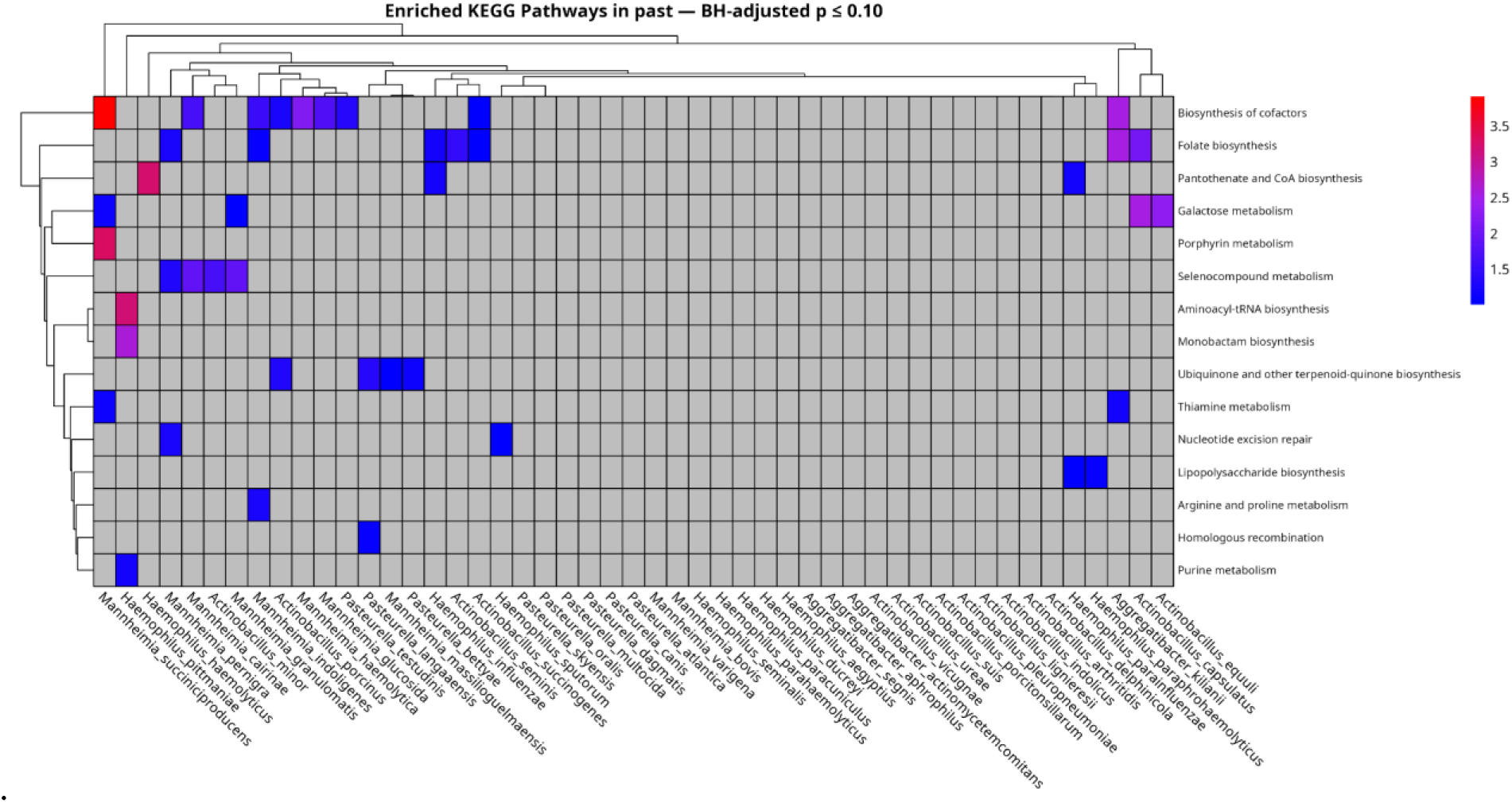
Enriched KEGG pathways in Pasteurellaceae. Color scale denotes -log10 adjusted p-values.

## Discussion

In this study, we investigated the distribution, diversity, and functional context of DUES across Pasteurellaceae and Neisseriaceae to understand how these motifs are shaped by evolutionary and functional constraints. Building on previous research and leveraging the large number of genome sequences now available through advances in sequencing technology, this study provides a comprehensive comparative analysis of DUES across Pasteurellaceae and Neisseriaceae. By systematically scanning for known and novel DUES dialects, we identified three previously undescribed DUES dialects, expanding the repertoire from 8 to 11 recognized dialects within Neisseriaceae. The genomic enrichment, which is even larger than for some well-established DUS in their respective genomes/hosts, and the similarity to previously reported DUS dialects strongly suggests they are genuine DUS dialects. The CTG core, which previously has been shown essential for transformation in assays using single transversion mutants of DUS, are conserved in the three new dialects. We have named the 10-mers with CTG in positions 6-8 according to convention (6) **GCCGTCTGCA** for **aniDUS** from of *N. animalis*, **GCGGCCTGTA** for **vitDUS** from *V. massiliensis* and **GCGTCCTGTA** for **vit3DUS** from *V. stercoraria*. The three respective genomes also encode the DUS receptor ComP solidifying the functional association between genomic DUS enrichment and DUS-specific DNA uptake.

Analyses of coding sequence context demonstrated that DUES are preferentially positioned in reading frames with lower biosynthetic energy costs, reinforcing the hypothesis that selection acts to minimize metabolic burden. Findlay and Redfield (11) showed significant USS bias towards the most common reading frames for the USS encoded amino acids alluding to translatability as selective pressure. They also found that USS preferentially were located in poorly conserved genes and in positions with poorly conserved amino acids. This does not fully align with previous results showing that DUS are enriched in the core genome, which generally encompass much more conserved proteins than those in the accessory (12). Indeed, we here find enrichment in GO and KEGG categories which are among the most conserved processes in nature, such as those involved in essential functions, replication, transcription, translation, cell wall synthesis, DNA repair and ATP hydrolysis. Many of these proteins are of considerable size and may have less conserved domains in which DUES potentially can reside without interfering with function. Deeper and systematic analysis of protein domains with DUES encoded amino acids and protein functions are warranted to further explore this potential type of selection. Taken together, it seems clear that DUES have evolved in CDS under strict Darwinian selective barriers. Furthermore, about half of all DUES were located within predicted transcriptional terminators, with inverted-repeat DUES showing even stronger localization (∼75% in Neisseriaceae and ∼90% in Pasteurellaceae), suggesting an additional level of structural and regulatory integration. This goes to show that the functioning of DUES inside transcriptional terminators is a near universally occurring phenomenon in these two families (5,9).

These observations beg the question as to why these genomes are so DUES enriched, even in CDS, when negative selection clearly influences their genomic placements. One model proposes that molecular drive from the DNA turnover mechanism itself causes DUES enrichment, given the selective DNA binding specificity by the DUES receptors ComP and ComN and considerable time. This model argues that DUES enrichment is a blind mechanism which saturates the genome with DUES in available and non-intrusive genomic space (26,27).; there is even translational space which DUES can occupy. It remains an open question however, if it is possible to evolve DUES enrichment across Pasteurellaceae and Neisseriaceae from molecular drive alone, that is, without any positive selection able to compensate for the evident negative selective pressures. In our opinion it remains highly questionable how sequence specificity could evolve in these families when they continuously would face the risk of being outcompeted into evolutionary oblivion by non-specific DNA uptake variants. DNA uptake sequence specificity is no longer a phenomenon confined to a few human pathogens, but is here identified and described in a wide array of bacteria living in very different environments. DNA uptake sequence specificity therefore seems a trait under positive selection. Finding different adaptations in diverse gram positive and gram negative species which bias DNA uptake to involve homologous DNA (28), supports a model where uptake and recombination evolved to mediate allelic reshuffling by homologous recombination (HR). Homologous recombination for allelic reshuffling is an ubiquitous feature of life and has been under such strong positive selection that it was maintained even when life evolved into meiosis-dependent sexual reproduction and multicellularity (29).

Functional annotation revealed that while the two families Neisseriaceae and Pasteurellaceae share a large proportion of GO terms overall, the overlap among enriched terms was markedly smaller, with Neisseriaceae showing a stronger bias toward genome maintenance processes such as “DNA repair”, “replication”, and basic cellular functions. KEGG pathway enrichment further supported this pattern, highlighting strong signals for “aminoacyl-tRNA biosynthesis” and “DNA repair” pathways in Neisseriaceae, but no corresponding enrichment in Pasteurellaceae. This signal of enrichment in genome maintenance functions in Neisseriaceae extends beyond the genus Neisseria, with members from the genera Kingella, Eikenella and Alysiella also showing some degree of enrichment in genome maintenance-related GO terms or pathways.

Relative to the Neisseriaceae, the Pasteurellaceae displayed narrower but distinct enrichments in homologous recombination and metabolic specializations. Although GO enrichment across the BP, MF, and CC ontologies generally shows weaker signals in Pasteurellaceae, a clear enrichment remains in GO terms related to genome maintenance. Notably, the biological process double-strand break repair via homologous recombination, the molecular function exodeoxyribonuclease V activity, and the cellular component exodeoxyribonuclease V complex (RecBCD) are all significantly enriched. RecBCD initiates the repair of DNA double-strand breaks through homologous recombination (30), and its consistent enrichment of related functions across all three GO ontologies represents a strong signal of selective importance in Pasteurellaceae. Thus, our results show a signal of enrichment of genome maintenance-related functions in both Neisseriaceae and Pasteruellaceae, even though the signal is stronger in Neisseriaceae. In this regard our observations align well with those of Davidsen et. al. (2004) (10), yet our results have better resolution which allowed the detection of differences between families also within the GMG category. Davidsen et al. (2004) (10) used a definition of genome maintenance genes (GMG) different from ours, applying the COG classification system (31,32) and a subclassification of group L (Replication, Recombination and Repair), whereas we used GO and KEGG terms in this study. Simultaneously, we do expect that many of the GMG genes are shared across Davidsen et al. (2004) (10) and this study, and thus some parallels can be drawn between the studies. It might also be argued that because GO terms are constantly being revised, that they are more up to date than the COG classification system (14,33). As such, this study serves as a corrective for DUES enrichment or at the least expands upon previous research. Davidsen et al. (2004) (10) used different mathematical and statistical models to measure DUES enrichment. The authors used a 4^th^ order Markov model to calculate the expected frequency of 9- and 10-mers and used the term relative abundance to refer to 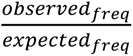 for k-mers. This can be a very accurate way of assessing DUES saturation within gene categories, but it depends on the veracity of the assumptions in the model used. This is a very different approach from our GO/KEGG enrichment analysis, which does not account for the number of DUES associated with a GO term. Findlay and Redfield (11) who re-analyzed the Davidsen *et al*. 2004 data (10) and included an additional *A. pleuropneumoniae* genome in their analysis and did not find the COG L+ category to have DUES densities higher than average. As we have discussed previously, the documented sequence specific transformability in the Actinobacillus are lower than that of *H. influenzae* by several orders of magnitude and this may impact how the enrichment analysis is interpreted (4).

Finally, finding DUES enrichment in the aminoacyl-tRNA biosynthesis genes in the Neisseriaceae aligns with the evolution of these fundamental enzymes where horizontal gene transfer has been highly significant (34). Altering translational fidelity and mistranslation have been shown to be potentially beneficial responses to environmental and cellular stress in different bacteria (35). DUES enrichment in these translation-linked categories could therefore further expand our understanding of transformation as means to respond to stress equivalent to stress-induced DNA repair.

One of the main strengths of this study is the large number of Pasteurellaceae and Neisseriaceae species analyzed. Earlier investigations of DUES enrichment in bacterial genomes were constrained by the limited availability of high-quality genomes, raising uncertainty about how broadly the findings applied to other bacteria with DUES. Beyond the greater number of species now included, this study also expands the ecological range considered. Earlier work focused exclusively on human associated microbes, both pathogens and commensals, whereas our dataset spans diverse hosts as well as free-living species. This broader scope is critical, since the genes conferring fitness are likely to differ depending on the ecological context in which a species exists. This is exemplified in *V. filiformis*, which was originally isolated from pond mud blackened by elemental sulfur deposits, an environment inhabited by sulfur-oxidizing bacteria such as *Beggiatoa* (36,37). *V. filiformis* is described as a non-sulfur-oxidizing, colourless bacterium with gliding motility. Unlike the other two *Vitreoscilla* species and all other Neisseriaceae, *V. filiformis* displays a distinct DUES enrichment profile in its very few enriched genes. It shows enrichment in fewer GO and KEGG categories, with the notable exceptions of pathways related to the sulfur cycle, bacterial chemotaxis (i.e. directional motility), selenocompound metabolism, and amino acid biosynthesis. These findings suggest that DUES enrichment in *V. filiformis* is closely tied to its high-sulfur environment and gliding motility appears to be a family-unique adaptation within the free-living *Vitreoscilla*. More broadly, the results indicate that enrichment of particular KEGG pathways and GO terms reflects selective pressures imposed by specific ecological contexts. Although H. influenzae and N. meningitidis inhabit overlapping niches in the human nasopharynx, they interact differently with their common hosts. *N. meningitidis* tends to adhere directly to epithelial cells (38) whereas *H. influenzae* is more associated with the respiratory mucosa and is more dependent on biofilm formation for colonization (39). It is therefore tempting to speculate that differential enrichment in DNA repair pathways, UvrABC and MMR in Neisseriaceae and RecBCD in Pasteruellaceae, to some extent reflect their divergent host interactions. For example, *N. menigitidis* which faces particularly strong oxidative bursts during blood-stream infections would be dependent on nucleotide excision repair by means of an immediate UvrABC response for survival through such dramatic and sudden bottlenecks. Survival of an *N. meningitidis* UvrA mutant has been shown reduced upon oxidative stress exposure (40). Furthermore, in being considered a lesser biofilm forming bacterium than than *H. influenzae* (41), *N. meningitidis* would benefit from tunable or mismatch repair (MMR) in order to mediate high rates of phase variation and adaptation in close contact with the host’s immune system. Indeed, null-MMR mutants are common in certain clinical strains of *N. meningitidis* suggestive of adaptive shifts in MMR activity which could be mediated by transformation of different MMR alleles (42). The RecBCD pathway is necessary for the more long-term adaptation from transformation, yet also the immediate rescue of double strand breaks from exposure to reactive oxygen and nitrogen species which would be essential for both species in the nasopharynx. Deeper analysis going from GO and KEGG categories to individual homologous proteins could shed light on potential links between bacterial lifestyles and DUES enrichment. If related groups of e.g. free-living bacteria from diverse environments were examined, it might reveal whether DUES are largely confined to more basal cellular functions like genome maintenance genes, or whether their distribution also reflects genes critical for adaptation to specific habitats.

This study has notable limitations. Although the number of species analyzed has increased, there is a strong overrepresentation of the genus *Neisseria*. Of all species examined, 38.1% (41/107) belong to this genus, accounting for 70.6% (41/58) of the Neisseriaceae species tested for enriched GO terms and KEGG pathways. As a result, shared and divergent signals between Neisseriaceae and Pasteurellaceae can be largely driven by *Neisseria*, and this must be considered when interpreting the results. The imbalance likely reflects a sampling bias: species closely associated with humans or other vertebrate hosts are more readily discovered as they occasionally may cause infections, whereas rarer taxa in more restricted ecological niches remain underrepresented. A fuller understanding of the evolutionary mechanisms underlying bacterial transformation and self-recognition requires comparative studies of distantly related groups from different ecological niches that have independently evolved similar systems. This underscores the importance of broader sampling efforts and ongoing large-scale genomics projects aimed at characterizing microbial diversity across diverse environments. Yet, the observed wide distribution of DUES across species living in highly different environments strongly suggests that DUES specificity itself is adaptive.

The methodological approach is also limited by this sampling bias. EggNOG-mapper annotates genes by assigning GO and KO terms based on the closest orthologous sequence in its database. For newly described species or those distantly related to well-characterized taxa, this often leaves many genes unannotated. An alternative is to use motif-based tools, such as HMMER2GO (github.com/sestaton/HMMER2GO?tab=readme-ov-file), which assign annotations based on sequence similarity rather than strict orthology. Additionally, studies have shown that certain genes and functional groups carry more DUES per gene, which may increase their likelihood of being transformed into related bacteria. This study did not assess that quantitative aspect (number of DUES pr. gene) as in previous studies (10,11), but future work should incorporate it and interpret enrichment signals with this in mind/by giving greater weight to genes with higher DUES counts relative to those with fewer.

